# Scoping Review of Maternal Monocytes and the Syncytiotrophoblast: Bi-Directional Communication in the Intervillous Space

**DOI:** 10.1101/2024.05.28.595900

**Authors:** Hannah Yankello, Caroline Smith, Kristin Daniel, Christina Megli, Elizabeth Wayne

**Author notes:** Authors contributed equally to this work (Caroline Smith, Kristin Daniel).

## Abstract

The relationship between maternal monocytes and the placenta is crucial for successful pregnancy. As existing reviews focus on the research on decidual macrophages or Hofbauer cells (fetal macrophages), we conducted a scoping review of literature studies examining the relationship between circulating maternal monocytes and the villous trophoblast. The goal of this review was to outline existing research within the field, disseminate further information, and to identify current knowledge gaps within the field. We searched the PubMed (MEDLINE) database for relevant articles published since 1995 that studied human tissue or cell lines. Our results showed clear trends in the type of monocyte and placental primary tissues and cell lines used. Further classification of these articles revealed four primary categories of monocyte-trophoblast interaction research: monocyte presence or recruitment to the placenta, monocyte phenotype in the intervillous space, monocyte-placental adhesion, and monocyte interaction with syncytiotrophoblast-derived extracellular vesicles (STBEVs). Although limited in scope and number, these studies implicate the importance of monocyte-trophoblast interactions, and their significant roles in maternal disease.

## 1. Introduction

A healthy pregnancy requires significant intercellular communication between maternal immune cells and the placenta^1^ to provide immunoprotection for the mother and immunotolerance for the fetus^2–4^. Much of this bi-directional communication occurs in the intervillous space, where circulating immune cells such as monocytes directly encounter the outermost layer of the placental villi called the syncytiotrophoblast (STB)^5^. This multinucleated cell^6^ releases significant amounts of immunomodulatory molecules^7^ and extracellular vesicles (EVs)^8^ believed to contribute to peripheral monocyte activation during pregnancy^4^, and a dysregulation of this cell-cell communication is associated with significant maternal and neonatal morbidities^9–12^.

Existing research studies and review papers on monocytes in pregnancy primarily focus on either tissue-resident decidual macrophages or Hofbauer cells (fetal macrophages). As such, we conducted a scoping review of literature characterizing maternal monocyte-syncytiotrophoblast interactions to highlight the known mechanisms of communication between these cells, identify current research methods used, and define existing gaps of knowledge within the field.

## 2. Methods

We selected to perform a scoping review of monocyte/trophoblast interactions as our primary goals for conducting this review closely aligned with three of the four reasons established for conducting Scoping reviews as outlined by Arksey and Malley^13^: our goal was to characterize the type, range, and depth of research occurring, summarize these research findings for increased knowledge dissemination, and outline current gaps of knowledge within the field^13^. With each step of this scoping review process, we closely followed the guidelines provided by Arksey and Malley^13^ as well as the checklist in the article “PRISMA Extension for Scoping Reviews (PRISMA-ScR): Checklist and Explanation”^14^. The overall methodology followed during this review process is illustrated in **Figure 1**.

**Figure 1:**
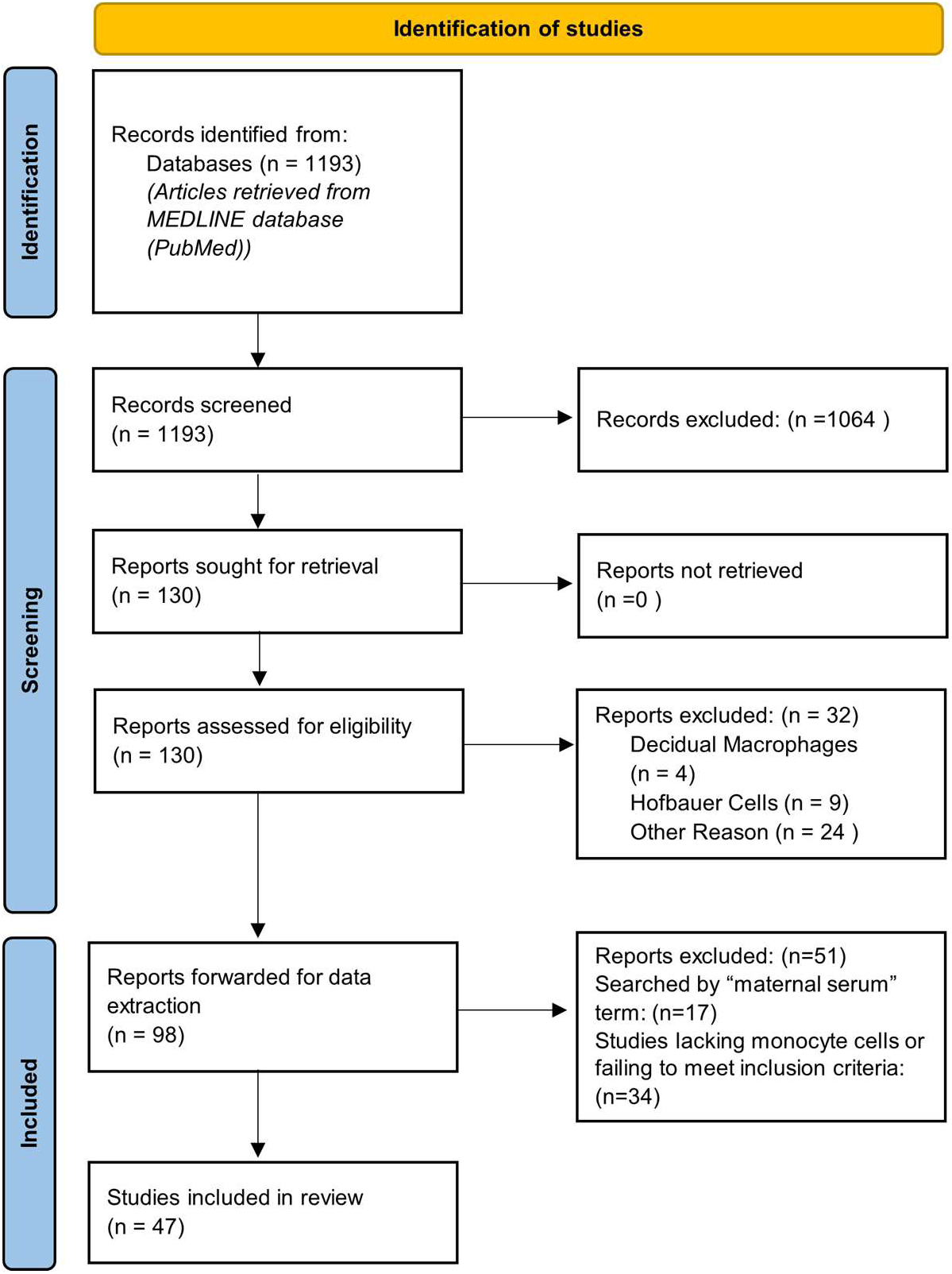
Methodology of the Scoping Review. The flow chart shows the step-by-step process for the collection and screening of articles included within this study. Of the initial 1,193 articles collected, 47 were included in this report. Chart adapted from PRISMA^14^.

### Database Search

We created a search string that was used to collect relevant literature within the PubMed (MEDLINE) database (**Supplemental Table 1**). The scope of the paper was to assess all studies characterizing interactions between circulating maternal monocytes and the villous trophoblast. This search string was composed of four primary groups. The first group identified all terms for the placenta as well as any cell lines known to model the villous trophoblast. The second group of terms specified that we were interested in articles pertaining to the intervillous space or maternal-fetal interface, the third group of terms primarily listed monocyte terms, and the remaining portion of the search string excluded irrelevant articles such as animal studies, review papers, or articles not published in English. Animal studies were excluded due to the significant differences between placental anatomy between species^15^. We limited the date of the publication of the study to between 1995-2023, as a few fundamental papers in this field were published before 2000, but many of the techniques or data from earlier records were outdated. The search string was finalized in August of 2023, and a total of 1,193 articles were collected from the database.

### Title-Abstract Screening

The collected articles underwent a dual-step screening process to determine whether the article should be included in the review. The initial screen included a manual screen of each article’s title and abstract to determine if the article met the inclusion and exclusion criteria outlined in **Table 1**. The screening was conducted using the Sysrev platform, and two individuals reviewed every article to minimize bias. Conflicts were resolved by a third independent reviewer. A total of 130 articles met the inclusion criteria after the completion of this initial screen. **Supplemental Figure 1** shows the reasons for exclusion of the collected articles at the title/abstract screening phase.

**Table 1:**
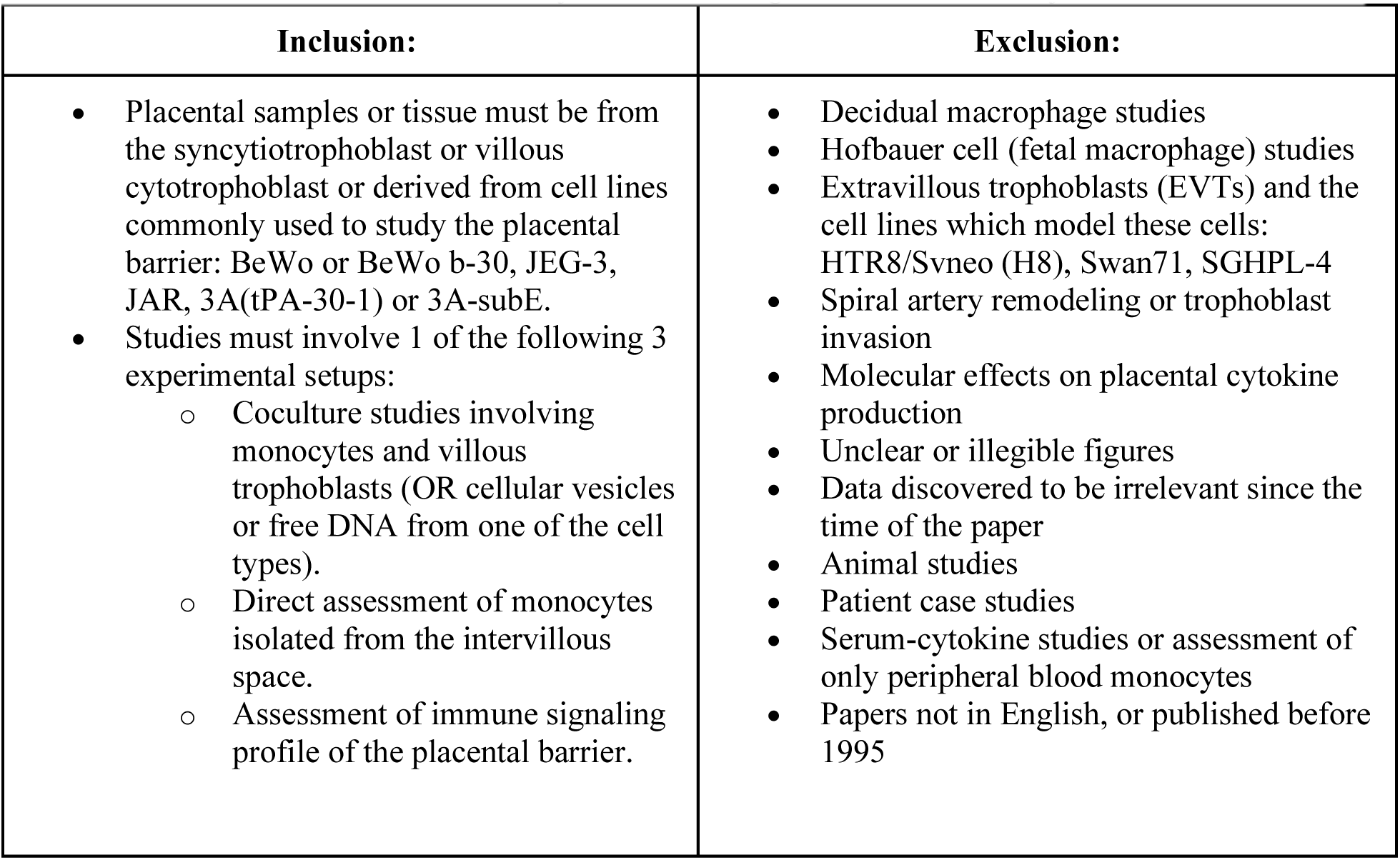
The inclusion and exclusion criteria for articles during the title-abstract and full-text screens.

### Full-Text Screening

After the initial screening, the 130 selected articles underwent a screen of the full-text of the article to ensure that the article met the inclusion criteria. This screening was again conducted using the Sysrev platform, and two individuals reviewed each article to minimize bias, with a third independent reviewer resolving any conflicts that arose. After this step of screening, 98 articles were selected to be included within the data extraction phase.

### Removal of studies without monocytes

Our initial search string and inclusion/exclusion criteria had been designed to collect reports of the placental secretion of known immunomodulatory factors into the maternal blood. As the addition of these studies would inherently introduce bias into the screening process (inclusion would depend upon the reviewers knowledge base of immunomodulators) and as our current search string was insufficient to collect articles relevant to placental cytokine production or steroid hormones, we decided to narrow the scope of our review to strictly studies that tested monocyte phenotype and function after interaction with trophoblasts or trophoblast signals. We ran our search string again without the “maternal serum” term on April 9, 2024, and removed all of the articles included after the full-text screen that were not located by that search. A total of 17 articles were removed.

### Data Extraction

After article screening, relevant data was extracted by a single user, based upon a survey of questions designed by input from all of the authors. The surveys questions used are outlined in **Supplemental Table 2**. Data was recorded and organized into spreadsheets. An additional 34 articles were excluded during this step. Many of the articles were not excluded in the previous step as they referenced “monocyte chemotactic protein” or other monocyte-related protein, or because their identification of monocytes in the tissue was found to not sufficiently delineate between maternal and fetal macrophages. The final number of reports included in the study was brought to a total of 47 articles. A list of included articles is recorded in **Supplemental Table 3**.

## 3. Results and Discussion

### 3.1 Overview of the field

Collected articles were published between 1995-August 2023 when the PubMed search was performed. **Figure 2A** demonstrates the number of included articles published each year within the included range of time. Overall, number of articles published per year did not appear to increase over time. The year which observed the highest amount of publications was 2021; however, this was likely due to publishing delays occurring in 2020 from the COVID-19 pandemic.

**Figure 2:**
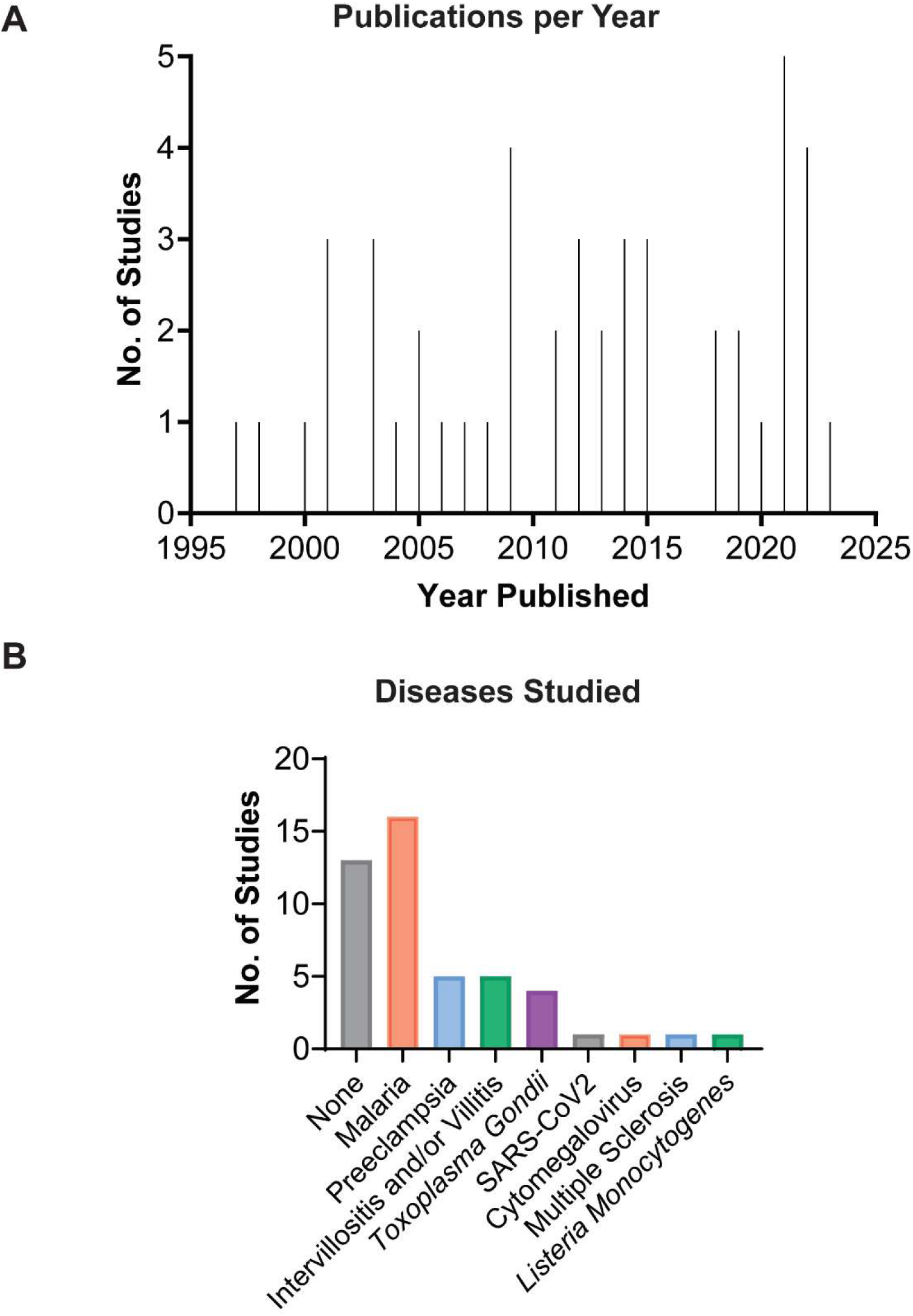
Overview of the field of research on the study of monocyte-trophoblast interactions in the intervillous space. A) Number of included articles published per year B) Number of studies examining specific maternal conditions

We then classified the different maternal disease conditions examined within the articles (**Figure 2B**). 34% of the articles studied malarial infections (either *Plasmodium Falciparum* or *Plasmodium Vivax*) infections. Unexpectedly, this number of articles was higher than the number of articles studying healthy pregnancy (27.7%). Overall, the collected articles examined only a limited number of maternal conditions, with five or fewer articles studying preeclampsia, intervillositis and/or villitis, and *toxoplasma gondii*. SARS-CoV2, cytomegalovirus, multiple sclerosis, and *listeria monocytogenes* were respectively studied by one article each. Of note, the intervillositis category included articles only discussing this condition. Other articles did observe intervillositis in the placenta as an effect of specific disease conditions and are discussed in section 3.5. In total, this graph highlights the need for further investigation into how maternal conditions dysregulate the monocyte-trophoblast relationship.

### 3.2 Placental Tissue Sources

We next outlined the sources and models of placental tissue used in each of the studies (**Figure 3**). The human placenta has three trophoblast subtypes, the cytotrophoblasts, the syncytiotrophoblast (STB), and the extravillous trophoblasts, each with a different function^16^. The human syncytiotrophoblast is generated from the fusion of underlying villous cytotrophoblast cells. It comprises the outermost barrier of the placenta and facilitates gas and nutrient exchange between the maternal and fetal circulations^17^.

**Figure 3:**
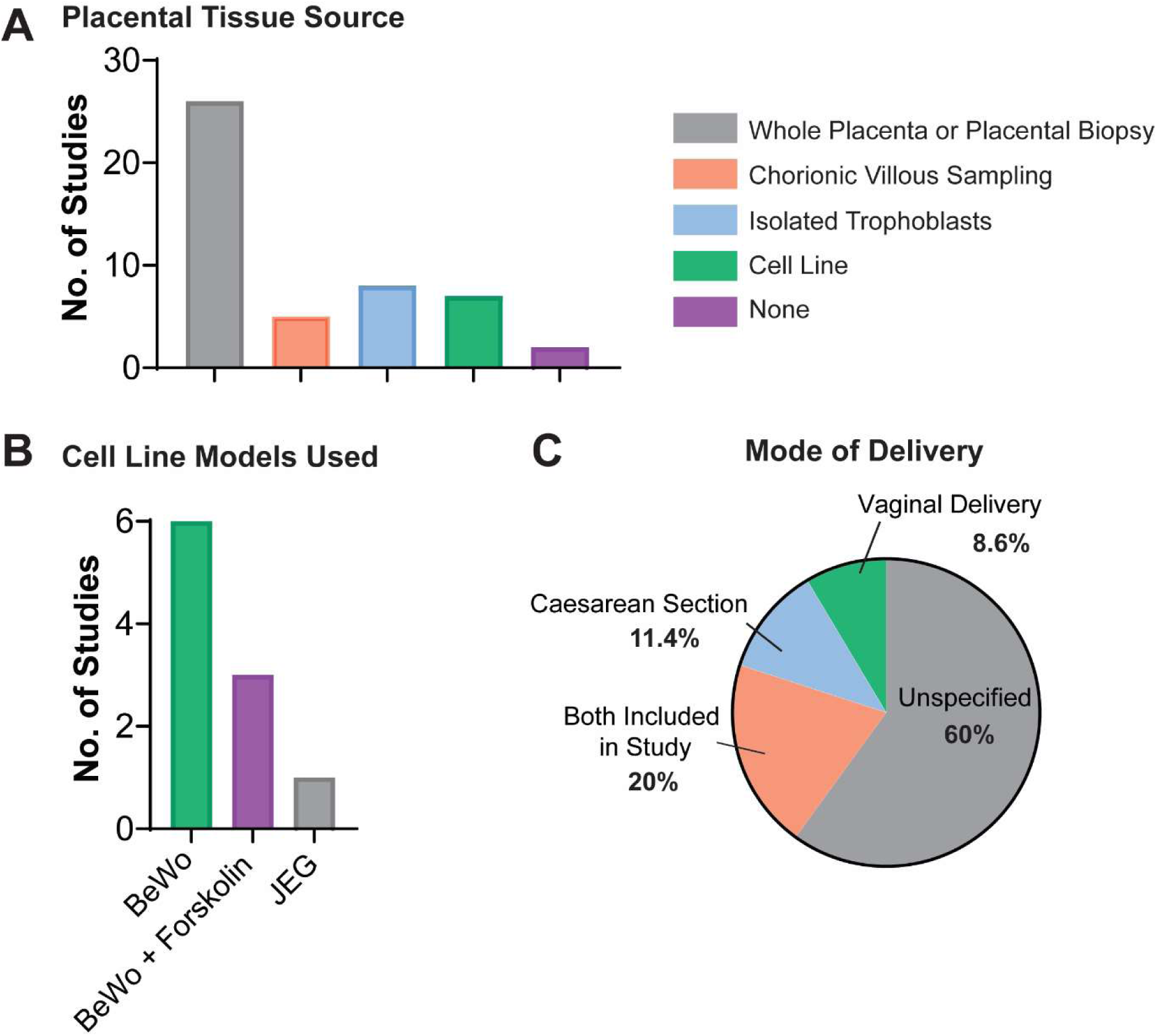
Overview of the types of villous trophoblast tissues and models used in the study of monocyte-trophoblast interactions. A) Frequency of placental tissue type used in the collected articles. B) The number of studies using choriocarcinoma cell lines and their respective types. C) Modes of delivery recorded in each of the studies

The vast majority of the articles obtained either used whole placentas or full-thickness placental biopsies for their studies (**Figure 3A**). These studies typically conducted immunostaining on the placenta to examine monocyte presence within the intervillous space or collected the intervillous blood on the maternal side of the placenta. Additionally, a couple of studies perfused whole placental lobules for the isolation of extracellular vesicles^8, 18^.

The remaining studies using placental tissue either performed chorionic villus sampling, isolated trophoblasts from tissue, or used an immortalized cell line, with one study using both chorionic villi and isolated trophoblasts^7^. Chorionic villous tissue is comprised of several different cellular populations^6^, and isolated trophoblast cells typically possess a short lifespan ex vivo^19, 20^. Consequently, the most widely used cell line models of the villous trophoblast are choriocarcinoma cell lines. Six of the reports used the BeWo cell line as a model of the villous trophoblast (**Figure 3B**). An additional three reported the use of forskolin-treated BeWo cells. Forskolin is a well-characterized agent which induces syncytialization of the BeWo cell line, making it the most significantly available model to study placental barrier function^21^. One additional studied used JEG cells^7^, another choriocarcinoma line well-documented in literature. As choriocarcinoma cells cannot fully recapitulate healthy trophoblast cells, the development of an immortalized trophoblast stem cell lines and organoids is an active area of research^22–26^. As these models are new, none were discovered within this literature analysis to examine monocyte-trophoblast interactions.

Of note, two articles did not use placental tissue in their study (**Figure 3A**). One of these articles tested a hormone secreted by the STB on human monocytes at the peripheral concentration levels observed in pregnancy^27^. The other article studied the phenotype of uterine vein blood collected during Caesarean section prior to the uterotomy and disruption of uteroplacental circulation^9^.

The gestational ages of the primary tissue used in the studies were not consistently recorded within each of the studies. What is recorded regarding these gestational ages is outlined in **Supplemental Table 3**. As expected, a significant portion of the primary tissues studied were derived from term placentas (>37 weeks gestation). Some articles included the average gestational age and standard deviation of the articles, while others simply classified the samples as term placentas. The exact percentage of studies using term placentas could not be calculated, as several articles used tissue from live deliveries, but did not specify the gestational age of the tissue. Based on the reporting available, four studies used tissue between 6-13 weeks gestation, and an additional four studies included tissue 14-28 weeks gestation. While these trends were expected and obvious, it highlights the significance gap of knowledge that exists for placental intercellular communication in the first and second trimesters.

Sixty percent of the thirty-five studies sourcing trophoblast tissue from live deliveries did not specify the mode of delivery (**Figure 3C**). A portion of the studies either only included vaginal deliveries (3 articles, 8.6%) or only included placentas from Caesarean deliveries (4 studies, 11.4%). Of note, all the studies limiting placental source to Caesarean delivery also excluded labored placentas. The remaining twenty percent of the studies (seven total) noted that both Caesarean and vaginal deliveries were included in the study, but only two of those studies examined labor as a variable within the investigation. Both of these studies demonstrated that human labor induces a pro-inflammatory state^28, 29^, and as such, future work should delineate tissue samples based upon labor or quiescence to minimize experimental variability.

### 3.3 Monocyte Sources

After placental tissue sources, we examined the types of monocytes used in the studies as well as their sources (**Figure 4**). As a significant portion of the studies did not isolate the monocytes from the tissue, but quantified monocyte presence and phenotype by immunostaining (27.7%) (**Figure 4A**). Just over half of the studies (53.2%) exclusively used primary monocytes isolated from blood, and one study used both primary monocytes and THP-1 cells. The THP-1 monocytic leukemia cell line was the only cell line recorded, and 17% of the studies used this as their sole source of monocyte cells. Compared to human primary monocytes, these cells possess a relatively quick doubling time, are easy to culture, and can be purchased at a relatively low cost^30, 31^. The behavior of this cell line has been extensively documented in literature as being a robust cellular model for human monocyte studies^32^. Nevertheless, as the THP-1 cell line is a cancer cell line and is derived from a one year old male^30, 31^, their reliability in the study of maternal health is often questioned. As such, most studies rely heavily on the isolation of primary human monocytes.

**Figure 4:**
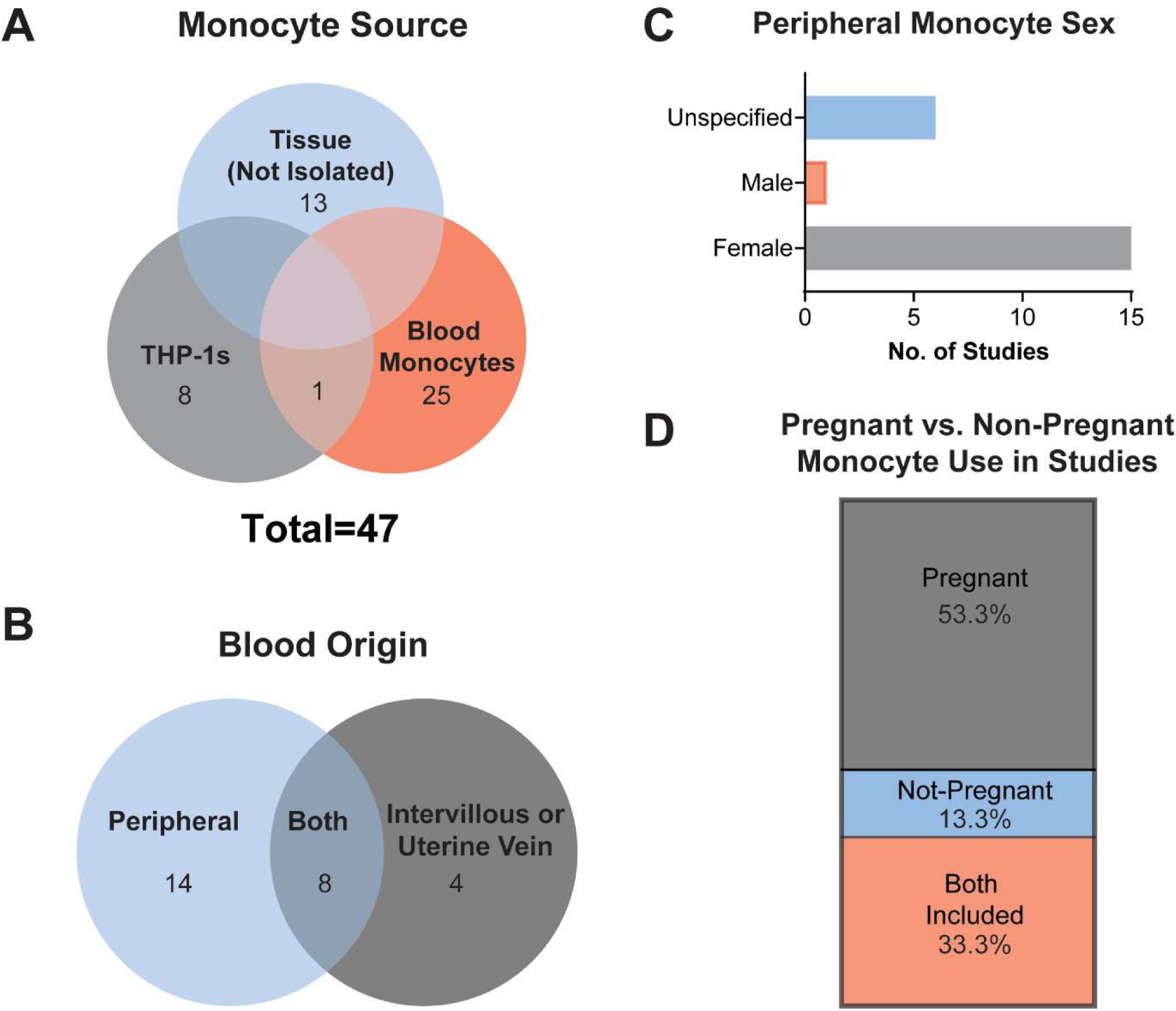
Overview of the sources and types of monocytes used in each of the included articles. A) A categorization of monocyte types in studies B) Sources of maternal blood monocytes. C) A characterization of peripheral monocyte sex utilized for relevant studies. D) Characterization of the pregnancy status of female peripheral monocytes used.

Both peripheral blood as well as blood from the uteroplacental circulation were examined within the collected articles. Maternal blood collected from the uteroplacental circulation was predominantly drawn from the intervillous space after delivery. One article collected blood from uterine veins during Caesarean sections just prior to the uterotomy, and as such, the blood collected had just exited the intervillous space^9^. Eight studies collected blood from both the peripheral and uteroplacental circulation of the same set of donors to compare the effect of the placenta on monocyte phenotype and recruitment (**Figure 4B**).

Increasing evidence demonstrates that biological sex can significantly impact innate immune responses^33^. Many of the genes encoding for innate immune responses exist on the human X chromosome^34^, and sex hormones have been demonstrated to modulate monocyte polarization^35, 36^. Consequently, we examined whether or not the peripheral blood monocytes used in the studies were from male or female patients. Most (68.2%) of all the studies using peripheral blood monocytes exclusively used female monocytes, while another six studies did not specify the sex of the monocytes, and one study used only male monocytes. Future work should emphasize the use of female monocytes in pregnancy studies, to accurately reflect monocyte state when it interacts with the syncytiotrophoblast.

Finally, as healthy pregnancy has been demonstrated to induce a pro-inflammatory response in the circulating monocytes^4^, we examined whether the reports specified if the female peripheral monocytes used were from pregnant donors. Just over half of these studies (53.3%) did recruit female peripheral monocytes, and another third of these studies used both pregnant and non-pregnant donors within the same study. The remaining 13.3% of the studies used non-pregnant donor monocytes. As the collection of pregnant monocytes is based on donor availability, it is likely that the sex and pregnancy status of the peripheral monocytes used was determined based upon availability as well as project timelines and resources.

### 3.4 Characterization of monocyte-trophoblast interactions

After examining the tissue sources used to study monocyte-trophoblast interactions, we categorized the types of interactions examined within each of the studies (**Figure 5**). Notably, several studies studied more than one type of interaction. Nearly half of the studies (24/47) observed monocyte presence in the placenta or studied monocyte recruitment to placental signals. Nine out of the 47 studies examined monocyte adhesion to villous trophoblasts and factors influencing this adhesion. Seven of the studies studied the phenotype of monocytes in the intervillous space, and five studies quantified the effect of syncytiotrophoblast-derived extracellular vesicles on monocyte phenotype and function. The remaining articles either observed the effects monocytes exerted on trophoblast function or described the impact of other secreted factors on monocytes. Each of these interactions is discussed in depth in the subsequent sections.

**Figure 5:**
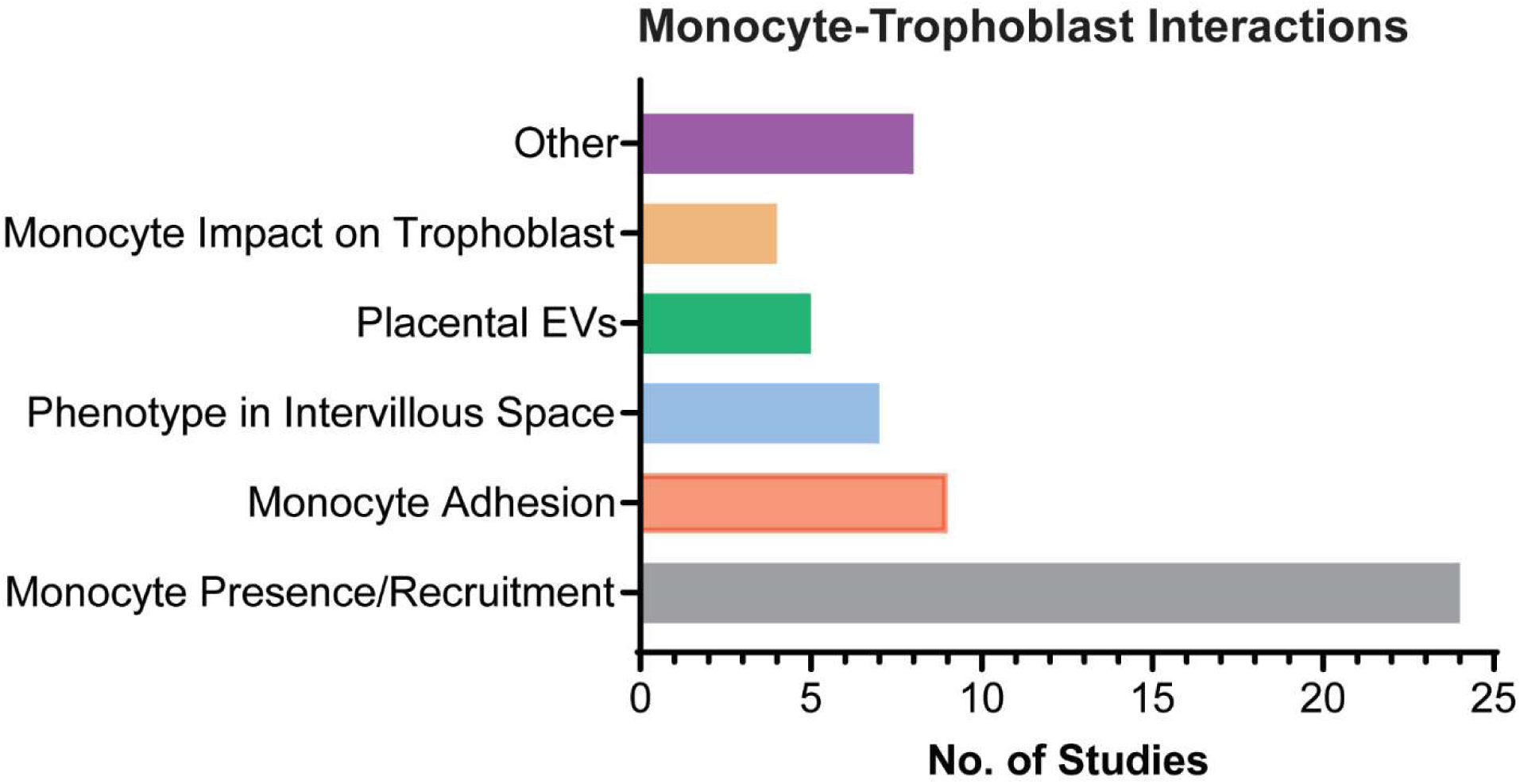
Monocyte-trophoblast interactions observed in the included articles. Several articles were included in more than one category.

### 3.5 Monocyte Presence and Recruitment in the Intervillous Space

As noted above, the highest percentage of the studies assessed monocyte presence or recruitment to the intervillous space of the placenta. Twenty-four articles total were included in this category. The experimental techniques for assessing monocyte presence or recruitment are shown in **Figure 6A**. Half of the articles surveyed used immunostaining with the CD68 macrophage marker to detect monocyte presence within the intervillous space, while another 25% (6 articles) used flow cytometry staining for the CD14 marker to detect monocytes. The remainder of the articles either used unspecified microscopic techniques (16.7%) or transwell migration assays (8.3%).

**Figure 6:**
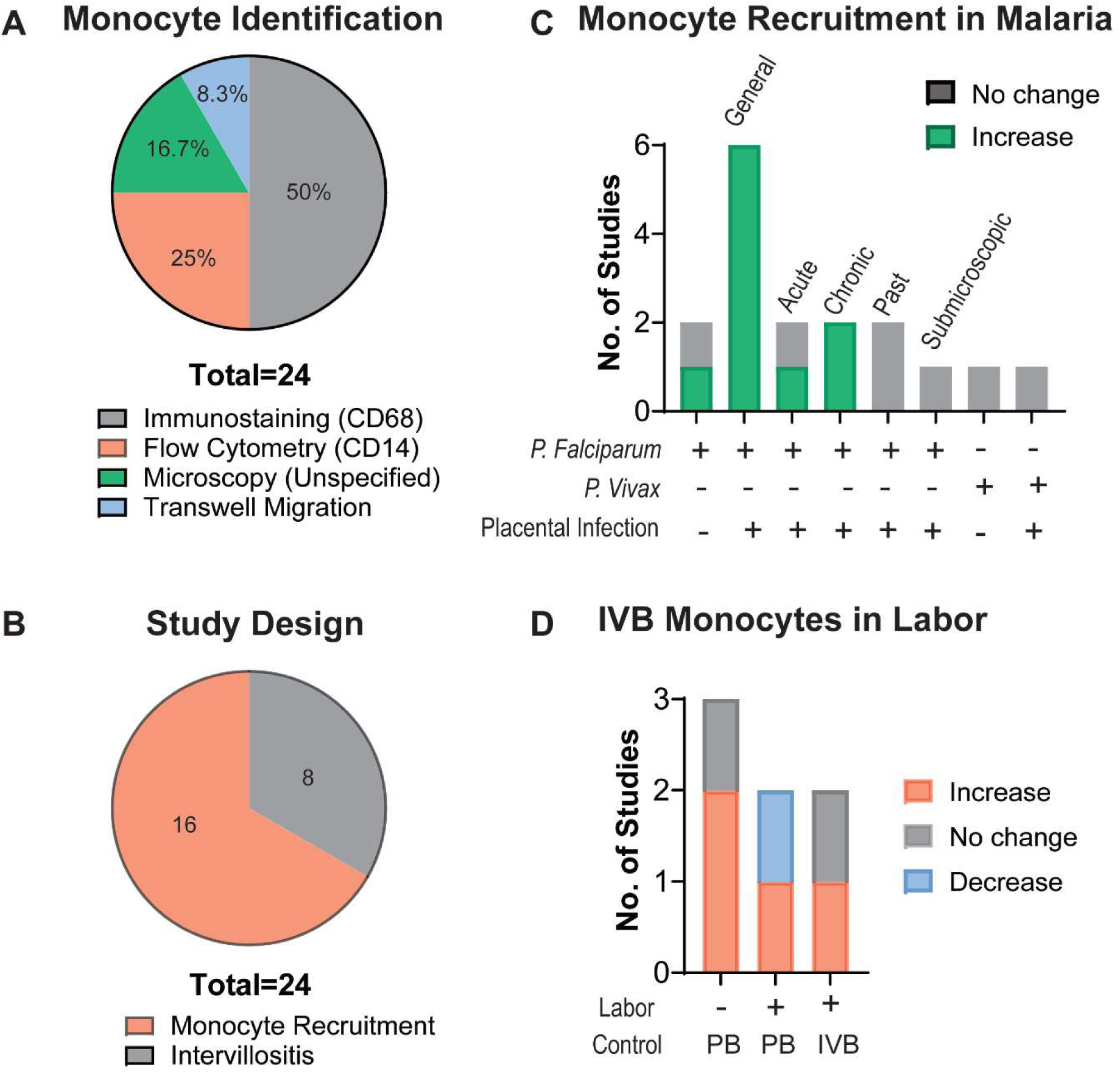
Monocyte presence/recruitment to the placenta. A) A categorization of the 24 studies included in this subset based upon the technique and surface marker used to detect monocyte recruitment. B) The study design of each of the 24 articles C) The changes and number of studies observing monocyte recruitment in malaria infections under different conditions. D) The changes and number of studies observing population changes of intervillous monocytes in labored and quiescent conditions compared to peripheral blood (PB) and intervillous blood (IVB) controls.

The objective of studies localizing monocytes to the placenta can be broadly categorized into the study of intervillositis (8 studies), or studies on monocyte recruitment in conditions such as placental malaria (where *P. Falciparum* parasites accumulate in the intervillous space, 9 studies), preeclampsia (1 study), and healthy gestation (6 studies) (**Figure 6B**).

The first group of studies broadly observed the presence of intervillositis in the placenta under different conditions, which included chronic villitis, malaria, and SARS-CoV-2. Intervillositis, also known as massive chronic intervillositis (MCI) or chronic histiocytic intervillositis (CHI) is a rare condition by which significant amounts of peripheral blood mononuclear cells are identified within the intervillous space of the placenta^37^. A few of these articles provided hypotheses as to why MCI might occur. The significant ICAM-1 expression in the syncytiotrophoblast in MCI was proposed as one possible cause^38^. As monocytes have the capacity to bind to the ICAM-1 ligand^39^, an increase in ICAM-1 expression could facilitate increased aggregation at the villi. Similarly, an increase in complement marker C4d was also observed on the microvillous membrane of the STB in MCI cases, implying an allogenic humoral rejection by recruited monocytes to trophoblast antigens^40^.

Several studies quantified monocyte recruitment to placental signals in *P. Falciparum* infections (**Figure 6 C**). Several noteworthy trends were observed. Six different studies consistently demonstrated that placental infection with the *Plasmodium Falciparum* strain of malaria increased monocyte recruitment to the placenta. When the subsets of these placental infection cases were examined, chronic placental infections alone demonstrated the highest monocyte presence. In contrast, past infections, submicroscopic placental infections (those only detectable by qPCR and not microscopy), and *Plasmodium Vivax* malarial infections did not affect monocyte recruitment to the placenta, and study results conflicted as to whether acute placental infections increased recruitment^41, 42^. This difference may be accounted for based on the type of control used in each study: one study used an uninfected placenta from the same region as the infected samples^41^, while the other used a control placental from another region unaffected by malaria^42^. Results also conflicted as to whether patient infections lacking placental infection caused an increase in monocyte presence^43, 44^. However, the differences in results may be due to differences in experimental methods used to detect placental infection^43, 44^.

Monocyte infiltration into the intervillous space in cases of *P. Falciparum* infection positively correlated with placental mRNA expression for chemokines MIP-1α, MCP-1, I-309, and IL-8 as well as intervillous blood concentration of CXCL10. *P. Falciparum* parasite burden also positively correlated with monocyte presence within the intervillous space^45^. Together, these correlations reveal the role of *P. Falciparum* infections in modulating trophoblast induction of monocyte chemotaxis.

One article examined monocyte recruitment to the intervillous space in preeclampsia (PE) within three subtypes of the disease. The subtypes were categorized by the clustering of gene expression data: cluster 1 (PE placentas which appeared mostly normal), cluster 2 (placentas with vascular malperfusion), and cluster 3 (PE placentas with evidence if elevated immune response). Using cluster 1 placentas as the control, cluster 2 placentas exhibited decreased monocyte presence, while cluster 3 exhibited increased presence, demonstrating that disease presentation impacts monocyte chemotaxis^46^.

Studies on monocyte recruitment in healthy gestation examined the impact of placental factors (**Table 2**) or human labor on monocyte recruitment (**Figure 6D**). Thomas et. al published an article demonstrating the presence of novel maternal monocyte populations in the first trimester placenta and suggest that peripheral blood monocytes are the precursors of these cells^47^. In term placentas, study results conflicted as to whether monocyte presence is increased in the intervillous blood compared to peripheral blood in both labored and non-labored conditions. Results also conflicted as to whether or not monocyte presence in the intervillous blood increased during labor^5, 48^. However, the study that found no changes between the non-labor and labor groups did observe an overall increase in monocyte presence, but the change was not significant^48^.

**Table 2:** Factors Affecting Monocyte Recruitment.

| **Article:** | **Trophoblast Stimulus:** | **Downstream effect:** |
| --- | --- | --- |
| Dallagi et al (2015)<sup>49</sup> | Leukemia Inhibitory Factor (LIF) treatment | Increase in the chemotactic potential of BeWo conditioned media (statistics not calculated) |
| Chen et al (2010)<sup>50</sup> | Lysophosphatidic acid (LPA) treatment | Increased chemotaxis as a result of GRO-α and MCP-1 induction. |

In total, the articles examining monocyte presence or recruitment in the intervillous space predominantly focus on disease conditions (**Figure 6A**). However, the signals and ability for monocytes to be recruited towards a healthy syncytiotrophoblast are yet uncharacterized. Further research should investigate the driving forces behind monocyte migration to the intervillous space, and their physiological role in successful pregnancy.

### 3.6 Monocyte Phenotype in the Intervillous Space

Seven articles collected in our search measured monocyte phenotype within the intervillous space. Of the seven articles, six examined monocytes in the third-trimester placenta while one examined monocyte phenotype in the first-trimester placenta **Figure 7A**. The first trimester study by Thomas et al identified novel populations of maternal monocytes and macrophages within the intervillous space, called PAMM1 cells (placental associated maternal macrophages)^47^. These PAMM1 cells can be further classified into two primary subtypes: PAMM1a cells exist as macrophages (FOLR2^-^CD9^hi^CCR2^lo/int^), while PAMM1b cells are true monocytes (FOLR12^-^ CD9^int^CCR2^+^) similar in phenotype to classical monocytes. When the cytokine secretion of these two populations of cells were compared, PAMM1b cells secreted greater amounts of pro-inflammatory cytokines compared to PAMM1a cells, indicative of their undifferentiated state^47^. This novel study provides a foundation for future research into maternal monocyte populations within the intervillous space.

**Figure 7:**
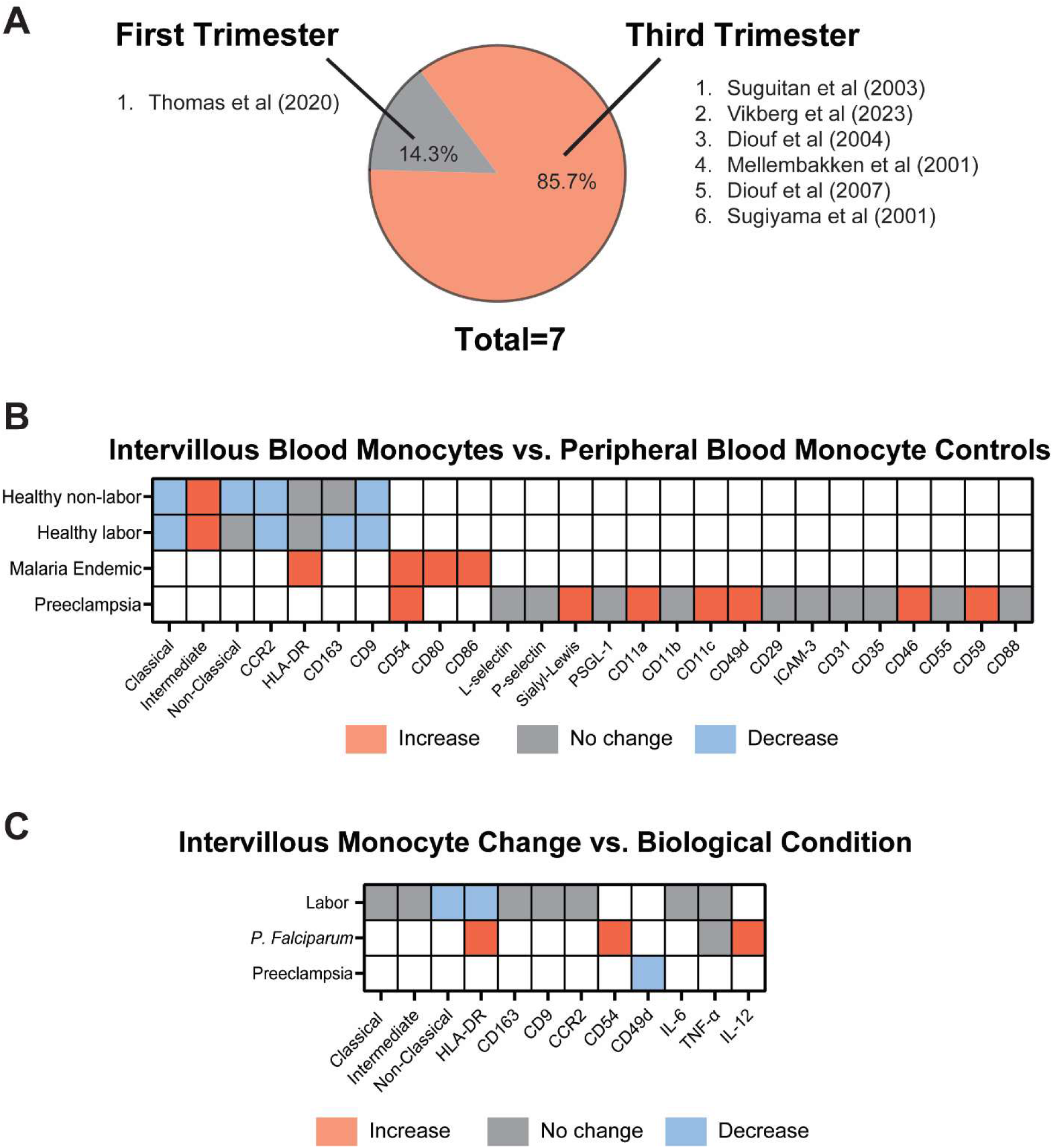
The phenotype of monocytes in the intervillous space. A) A breakdown of the 7 studies within this subset based upon pregnancy trimester. B) Changes in surface marker expression observed between intervillous blood monocytes and peripheral blood monocytes under varying conditions. C) Phenotypic changes observed in intervillous blood monocytes in labor, malaria infection, or preeclampsia.

The remaining six studies examined the phenotype of third-trimester monocytes in the intervillous space or uterine veins at delivery, with one study demonstrating that altered production of TNF-α, IP-10, MCP-1, MIP-1α, and MIP-1β by intervillous white blood cells was correlated to the percentage of monocytes in the culture^41^. The remaining studies focused on surface receptor surface receptor changes occurring in intervillous blood monocytes as compared to peripheral blood of the same donors, or compared to an intervillous blood control (**Figure 7B,C**).

Recently, a study by Vikberg et al applied the gating scheme for PAMM1a cells developed by Thomas et al to monocyte populations in term intervillous blood. The PAMM1a gating scheme was selected due to the lower expression of CCR2 on the monocytes in the intervillous blood^5^. The study did not find evidence of a PAMM1a cell population in the third trimester intervillous blood^5^; however, as PAMM1a cells were characterized as adherent macrophages^47^, it is possible that the process of collecting blood from the intervillous space failed to capture this population.

When comparing intervillous blood monocytes to their peripheral blood counterparts, a general observation was noted. The diseased groups demonstrated that IVB monocytes had increased or no change in expression compared to their peripheral blood counterparts, while the healthy groups (excepting the intermediate) typically exhibited either no change or decreased surface marker presence. Vikberg et al. hypothesized that the increase in intermediate populations in intervillous blood vs. peripheral blood is likely correlated to the decrease in the classical monocyte markers CD9 and CCR2^5^.

CD54, also known as ICAM-1 was found to be upregulated in intervillous blood monocytes vs. peripheral monocytes in two separate studies. One study collected patient blood samples with and without *P. Falciparum* infection in a malaria endemic region, but did not outline whether the cells assessed for CD54 presented with a malaria infection or not^51^. The other study observed an upregulation of CD54 expression on intervillous monocytes in preeclampsia^9^. CD54 expression has been demonstrated to be upregulated on CD16+ monocyte populations (intermediate and non-classical monocyte subsets)^52^ and has specific roles in monocyte-T cell interactions^53^, highlighting again the influence of the placental barrier on immune responses in the intervillous space.

When changes in IVB monocyte markers were observed in different conditions (**Figure 7C**), only a couple of markers were repeated across different studies. HLA-DR expression in IVB monocytes was discovered to decreased in labor but increase in the context of *P. Falciparum* infection. HLA-DR is a well-characterized marker associated with the intermediate monocyte subset (CD14^hi^CD16^+)54^. Roles of the intermediate monocyte subset include pro-inflammatory cytokine secretion, antigen presentation, and T-cell stimulation^55, 56^. In the case of labor, the elevated cortisol levels in human labor are suggested to suppress HLA-DR expression in intervillous blood monocytes^5^ despite overall no change in IVB intermediate monocytes^5^. In contrast, *P. Falciparum* infection in the placenta elevated IVB monocyte expression of HLA-DR, which is hypothesized to be the result of the increased TNF-α and IFN-γ levels during malarial infection^51^.

### 3.7 Monocyte Adhesion to the Syncytiotrophoblast

Another relevant interaction between maternal monocytes and the syncytiotrophoblast observed in the collected studies was the adhesion of the maternal monocytes to the placental barrier. A total of nine articles observed this phenomenon. Four of these articles made general observations regarding the adhesion of maternal cells to the STB, but did not quantify the number of cells adhered or make comparisons between groups (**Figure 8A**). The most notable of these articles was the paper discussed above by Thomas et al published in 2020^47^, demonstrating that maternal monocytes adherent to the first-trimester syncytiotrophoblast differentiate into a novel population of maternal macrophages. These macrophages were called PAMM1a cells (placental associated maternal macrophages) and were characterized by the markers FOLR2^-^ CD9^hi^CCR2^lo/int^. Of significant note, these PAMM1as were observed to be present surrounding breaks in the syncytiotrophoblast, indicative of their potential function in tissue repair^47^.

**Figure 8:**
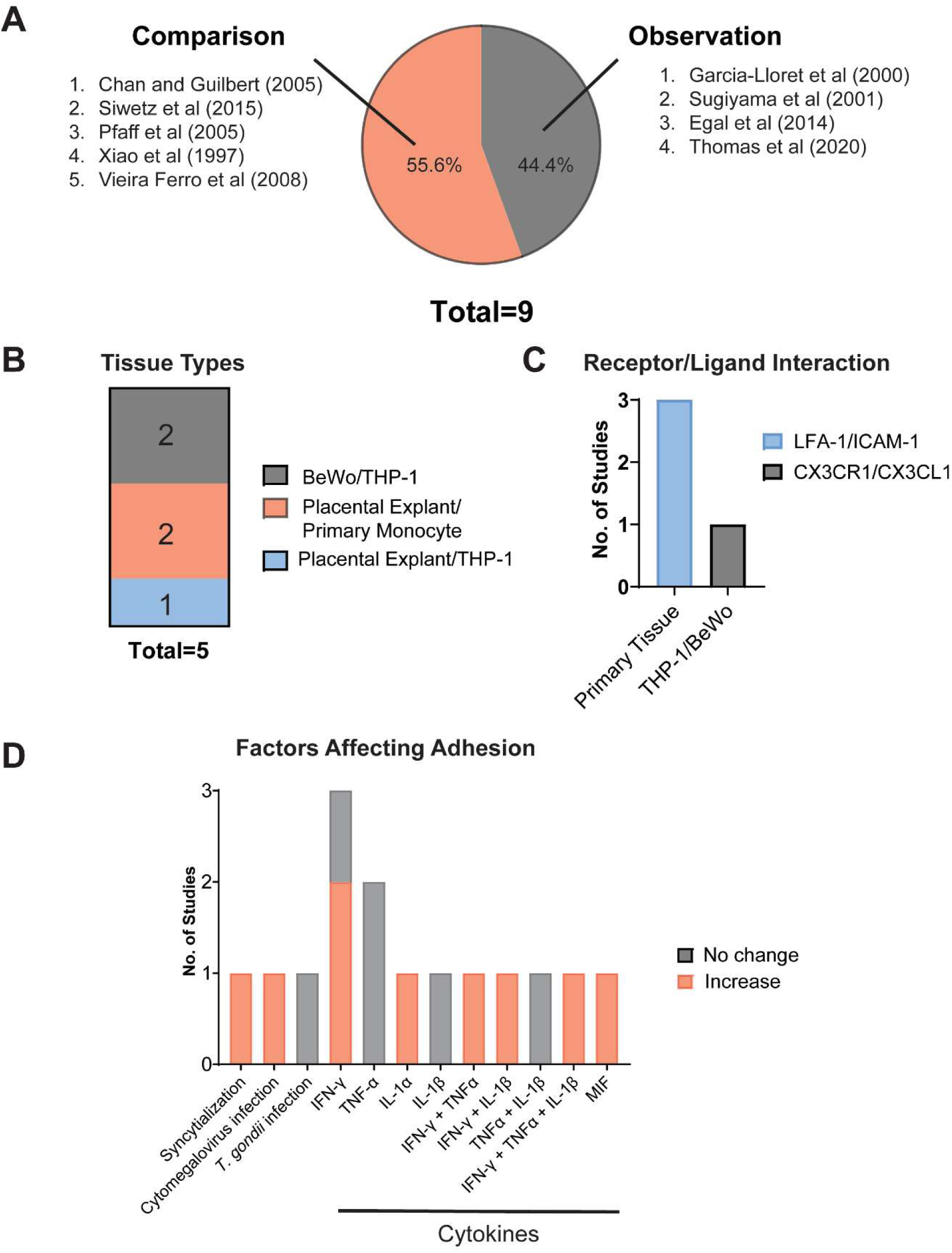
Monocyte adhesion to trophoblast tissue. A) A categorization of the 9 studies included in this subset based upon whether they simply observed monocyte adhesion or compared adhesion changes under specified conditions. B) The tissue types and pairings of monocytes and trophoblasts demonstrated to facilitate monocyte adhesion. C) The number receptor/ligand interactions quantified in both primary tissues and cell lines. D) Cellular factors impacting monocyte adhesion

The other five articles collected observed the changes in monocyte adhesion to placental tissue dependent upon changes in the cellular microenvironment. These experiments were designed with various pairs of cell lines and primary tissues **Figure 8B**. While THP-1 cells were observed to adhere to both BeWo cells^57, 58^ and placental explants^59^, none of the studies examined the adhesion of primary monocytes to BeWo cells. It is unclear whether this is due to experimental design, or because primary monocytes actually do not adhere to the BeWo cell line (experiments completed our lab indicate that the latter reason may be the case).

Several factors were demonstrated to increase monocyte-placental adhesion (**Figure 8D**). Trophoblast syncytialization^57^ or infection with cytomegalovirus^60^ were the cell specific factors identified. Other pro-inflammatory cytokines such as IFN-γ, IL-1α, and MIF were also implicated as upregulating monocyte-placental adhesion^59, 61^. Of note, IFN-γ alone was shown to increase monocyte adhesion in explant cultures^60, 61^, but failed to increase THP-1 adhesion to BeWo cells^58^. A similar observation was noted for IL-1^58, 61^. As BeWo cells are a choriocarcinoma cell line, it is expected that differences will be observed between those cells and primary tissue.

Lastly, four of the articles performed mechanistic studies to identify the ligand/receptor interactions present during monocyte-placental adhesion. Three demonstrated that monocytes adhere via the LFA-1 receptor to ICAM-1 on the syncytiotrophoblast^39, 60, 61^, with all the experiments completed in exclusively primary cells. The CX3CR1/CX3CL1 interaction was also implicated; however, these experiments were done entirely in cell lines^57^. To our knowledge, no studies have identified this interaction in primary tissue.

As many of these articles emphasized the role of syncytiotrophoblast ICAM-1 expression in monocyte adhesion^58, 61, 62^, it is implied that other diseases associated with increased ICAM-1 expression on the STB may also exhibit increased monocyte adhesion. Future work is needed both to categorize these cell-cell interactions, as well as to understand the downstream effects of this interaction.

### 3.8. Monocyte Interaction with Syncytiotrophoblast-Derived Extracellular Vesicles

Five of the collected articles focused solely on the interaction of syncytiotrophoblast-derived extracellular vesicles (STBEVs) with circulating immune cells and are listed in **Table 3**. Microscopic examination of the blood-exposed surface of the STB contains significant microvilli, which profusely release microvesicles or microparticles (100-1000 nm) into maternal circulation. The syncytiotrophoblast also possesses large numbers of multi-vesicular bodies, which can fuse with the plasma membrane to release exosomes (30-100 nm) into the bloodstream^63, 64^.

**Table 3:** A list of the articles that examined the influence of STBEVs on maternal monocytes, and their reports regarding vesicle binding and uptake.

| Article Cited: | STBEV Binding or Uptake by Monocytes |
| --- | --- |
| Messerli et. al<sup>65</sup> | • Not characterized |
| Southcombe et. al<sup>18</sup> | • 82% of monocytes bind to STBEVs 1 hr after treatment<br>• 90% of monocytes internalized STBEVs 20 hrs after treatment |
| Germain et. al<sup>8</sup> | • % of monocytes bound to STBEVs in bloodstream increases across the three trimesters of pregnancy. |
| Martire et. al<sup>66</sup> | • Not characterized |
| Bai et. al<sup>67</sup> | • >90% of monocytes had interacted with STB exosomes after 24 hrs incubation |

Germain et. al showed that these STBEVs can be isolated directed from maternal peripheral plasma, and that these particles in the bloodstream circulate both freely and bound to circulating monocytes^8^. The numbers of both free and bound STB microparticles in plasma increased over the progression of pregnancy^8^. As the cargo of extracellular vesicles reflects both the cell type and disease state, STBEVs hold promise as potential biomarkers for maternal diseases such as preeclampsia^68^.

Three primary methods were described to prepare harvested placental explants for the isolation of released EVs. **Figure 9A** illustrates number and type of techniques adapted by each of the five articles. The first technique was stationary incubation, whereby the placental explant was incubated in cell culture media at 37 degrees Celsius without disruption for a period of 24-72 hrs^18, 65–67^. As the intrauterine tissue microenvironment is hypoxic^69^, and tissue hypoxia has been demonstrated to increase EV production^70, 71^, some of these studies cultured the explants under low oxygen tension as well. **Figure 9B** shows the differences in length of incubation and oxygen tension for stationary-incubated placental explants.

**Figure 9:**
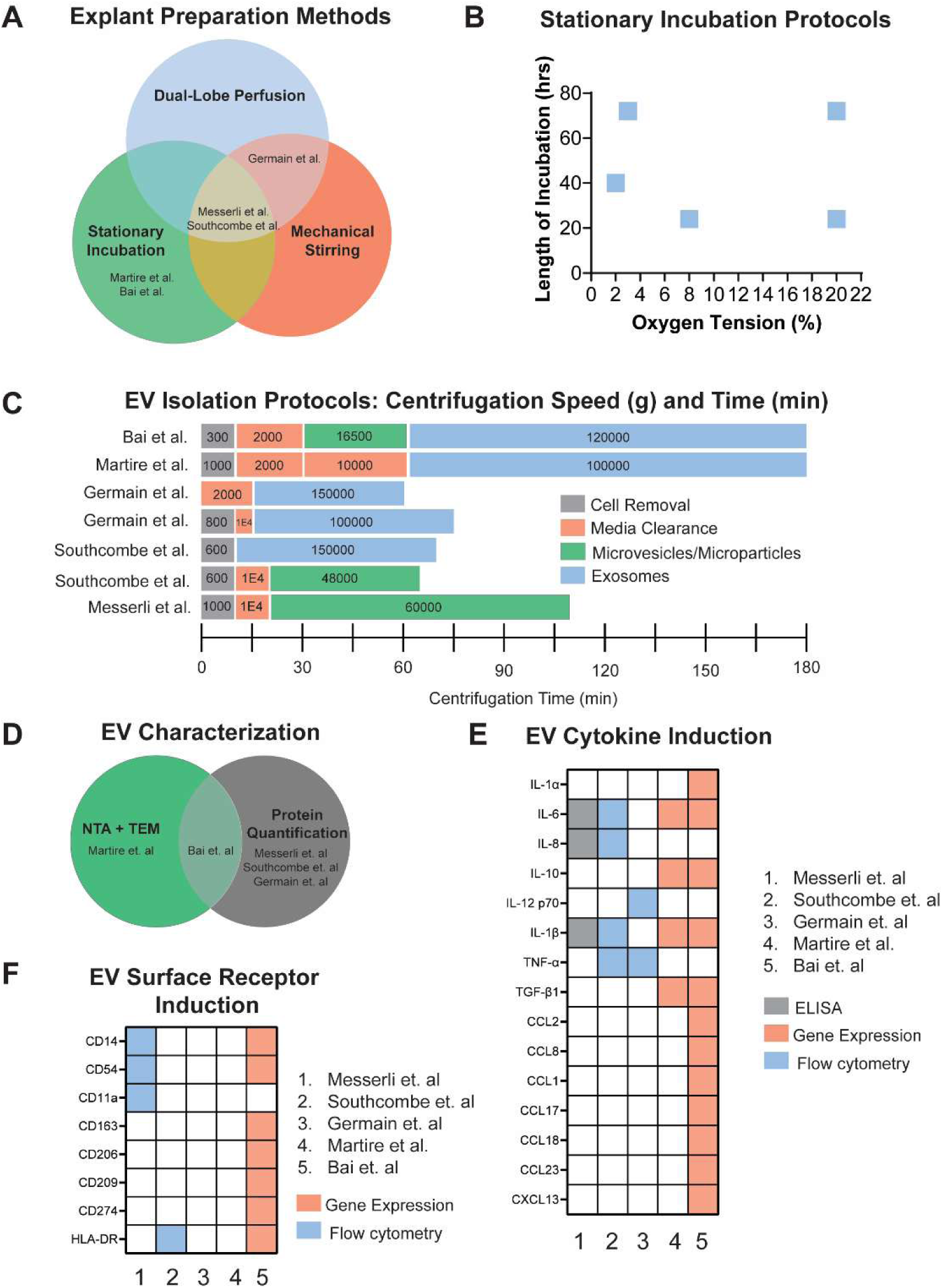
Monocyte interaction with syncytiotrophoblast-derived extracellular vesicles. A) Explant preparation methods used by the five studies for future EV isolation. B) Length of incubation time and oxygen tension parameters used in the stationary incubation protocols. C) Differential ultracentrifugation protocols for the isolation of EVs. D) EV characterization methods used in each of the studies. E) Monocyte cytokine production following STBEV interactions and the methods used to detect cytokine induction. F) Surface receptor changes in monocytes following interaction with STBEVs and the technique used to assess these changes.

The second method to prepare explants for EV isolation was mechanical stirring, where pieces of placental explant were mechanically stirred at 4 degrees Celsius in saline solution. The time of stirring varied between 30 minutes to overnight^8, 18, 65^. The final method, dual-lobe perfusion, involved the perfusion of the intervillous space of a whole placenta with specialized medium for a period of 20-30 minutes^8, 18, 65^. Of the three studies that compared EVs isolated from different methods^8, 18, 65^, all demonstrated that mechanically stirred EV preparations failed to induce cytokine production in monocytes and peripheral immune cells, in contrast with the other two methods. It is unclear whether this difference is actually due to the technique itself, or due to downstream isolation procedures, as in 2 out of three studies the mechanically-stirred EVs were then isolated via a different centrifugation regime compared to the perfusion or stationary EVs^8, 18^.

The differential ultracentrifugation protocols followed in each study are illustrated in **Figure 9C**. Each lab used a slightly different isolation protocol or protocols of different centrifugation times and speeds. Two of the protocols did not exceed 60,000g in for the final centrifuge step, and as such only isolated larger EVs.

EV characterization methods also varied between the papers. Three of the papers only quantified the protein content of the EVs, one paper used nanoparticle tracking analysis (NTA) and transmission electron microscopy (TEM) to record the size, concentration, and morphology of the EVs, and the final paper did all three characterizations (**Figure 9D**). Due to the differences in explant preparation, EV isolation, and EV characterization, it is difficult to make comparisons between the results obtained between the studies. As such, only overall characterization trends are discussed.

Two of the five articles demonstrated that of peripheral immune cells, monocytes are the dominant cell type that interacts with the EVs^18, 67^ (**Table 3**). Southcombe et al. demonstrated that within 1 hr of treating monocytes with EVs (derived by dual-lobe perfusion), 82% of the monocytes had EVs bound to their surface, and after a period of 20 hrs, 90% of monocytes had internalized the STBEVs^18^. Similar results obtained by Bai et al. revealed that after 24 hrs of incubation with trophoblast EVs (derived by stationary incubation), over 90% of monocytes were interacting with the vesicles^67^ (**Table 3**).

After incubation with STBEVs, monocytes were characterized based upon either gene or protein expression of cytokines and surface receptors. **Figure 9E** shows the cytokines characterized and method of characterization for each of the studies. The most characterized cytokines were interleukin-6 (IL-6) and interleukin-1β (IL-1β), both of which are pro-inflammatory^72^. Similarly, **Figure 9F** illustrates the surface receptors examined following monocyte incubation with trophoblast EVs and the method of characterization used. As surface markers examined in more than one study often used different measurement techniques, it is not possible to draw detailed analyses from the data.

### 3.9 Other impacts of the placenta on monocyte phenotype and behavior

Lastly, all other observed signaling from trophoblast cells to monocytes was recorded in **Table 4**. A few of these articles specified the secreted protein responsible for the monocyte response observed. The syncytiotrophoblast is a proliferative producer of cytokines^73^, hormones^74^, and cell-free fetal DNA^75^ and releases these components directly into circulation. Numerous articles have quantified the secreted cytokine and steroid hormone profiles of the placental barrier, but these articles were outside the scope of our search. The broad range of the articles below highlight the further need to investigate the effects of placental signaling on maternal monocytes.

**Table 4:**
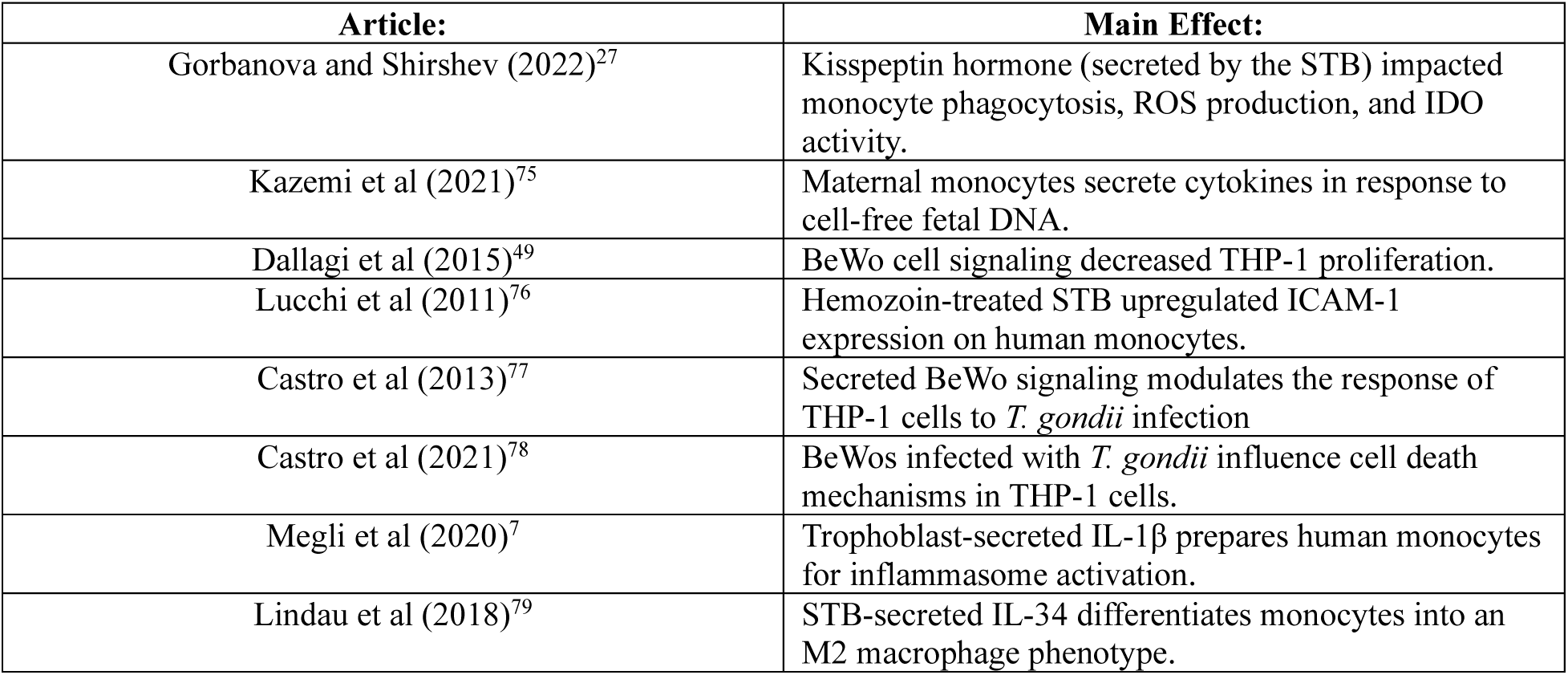
Other included articles which characterized monocyte-trophoblast interactions.

### 3.10 Monocyte impact on the placental barrier

A final query investigated in this collection of articles was whether monocyte signaling to the placental barrier affected its phenotype or barrier function. Only four of the articles collected analyzed this question. Two of the articles noted the impact of adherent monocytes on placental cell death. Chan and Guilbert observed that monocyte binding to healthy placental explants did not upregulate placental apoptosis^60^. However, monocytes adherent to placentas infected with cytomegalovirus did increase placental cell death. This result was found to be significant when compared to the levels of cell death occurring due to viral infection^60^. In contrast, another study found that adherent monocytes did increase cell death of synytiotrophoblasts, and this was determined to be a local effect driven by the secretion of TNF-α^39^. The differences in these results can be attributed to cellular conditions, as Garcia-Lloret et al pre-incubated monocytes with lipopolysaccharide (LPS) and the syncytiotrophoblast cells in the presence of IFN-γ to simulate placental inflammatory conditions^39^. As such, it is hypothesized that monocytes may enhance the detrimental effects of infection and inflammation in the placental barrier.

Another effect of monocytes on the placental barrier pertains to the amino acid uptake of trophoblast cells^80^. Supernatant from cocultures of monocytes and *P. Falciparum* infected erythrocytes were found to decrease the amino acid uptake of BeWo cells relative to uninfected supernatants. It was suggested that IL-1β may have an impact in this decrease, but this was discovered to not be the case^80^.

Finally, Rice et al examined the effects of both monocyte and macrophage-derived extracellular vesicles on placental cytokine production^81^. Of note, they discovered that monocyte EVs did not affect cytokine production by trophoblasts, but macrophage EVs did induce a significant increase in production^81^. As it has been demonstrated that some monocytes in the intervillous space differentiate into macrophages^47^, we hypothesize that this PAMM1a population may have a significant role in regulating syncytiotrophoblast secretion of cytokines via EV signaling.

## 4. Conclusions and Future Work

To our knowledge, this is the first scoping review conducted of monocyte communication with the syncytiotrophoblast. From the reports collected, the significant limitations regarding the depth and range of research conducted become apparent. As such, it was difficult to draw significant conclusions regarding the monocyte-trophoblast relationship. However, we broadly observed four different categories of monocyte-trophoblast communication tested: monocyte recruitment into the placenta, monocyte phenotype changes in the intervillous space, monocyte-placental adhesion, and monocyte interaction with STBEVs. Additional articles examined the impacts of other signaling molecules on monocyte functional behavior. Only a few articles implicated the effects of monocytes on placental explants, suggesting that monocytes are implicated in placental apoptosis and amino acid transport.

Given these findings, future research in this field should contain several factors: First, as emerging research demonstrates that labor induces a pro-inflammatory state^5^, research on placental explants should include the delivery method of the explants, as well as note for Caesarean deliveries whether the tissue had labored or not. Similar documentation should be expanded for the use of primary human monocytes, as biological sex impacts cellular immune responses^34^.

Second, given that increased recruitment of monocytes in the intervillous space can be associated with negative pregnancy outcomes^37^, the study of monocyte recruitment in the placenta should be expanded to other maternal disease beyond *P. Falciparum* to identify how placental chemotactic signaling changes within different contexts. These studies could provide valuable insight into novel biomarkers of disease.

Third, an expansion of the characterization between intervillous blood monocytes and peripheral blood monocytes should be conducted. Significant research focuses on changes in peripheral monocyte populations during maternal disease, but further insight into how the IVB and peripheral populations differ will assist in the accurate stratification of disease as well as overall placental health.

Lastly, future work should characterize the impact of circulating monocytes on the placental syncytiotrophoblast, as well as the impact of differentiated macrophage populations in the intervillous space. Due to the circulatory nature of human monocytes, an understanding of their impact on the placental can assist in the monitoring placental health.

