## Supplemental Material for "Scoping Review of Maternal Monocytes and the Syncytiotrophoblast: Bi-Directional Communication in the Intervillous Space"

**Supplemental Tables and Figures**

**Supplemental Table 1: Search String for MEDLINE (PubMed)**

***Note:*** *Article search was conducted on August 25, 2023*

| String Number: | Search Terms: | Number of Articles Identified: | Purpose of Search Terms: |
|---|---|---|---|
| #1 | villous trophoblast*[tiab] OR cytotrophoblast*[tiab] OR syncytiotrophoblast*[tiab] OR primary human trophoblast*[tiab] BeWo*[tiab] OR JEG-3*[tiab] OR 3A-subE*[tiab] OR 3A(tPA-30-1)*[tiab] OR JAR*[tiab] OR JAr*[tiab] OR trophoblast[MeSH] OR placenta[MeSH] | 79,054 | *Identify relevant placental terms or trophoblast cell lines.* |
| #2 | maternal-fetal interface*[tiab] OR feto-maternal interface*[tiab] OR intervillous space*[tiab] OR Maternal-Fetal Exchange[MeSH] | 32,598 | *Identify articles studying the maternal-fetal interface.* |
| #3 | Monocyte[MeSH] OR monocyte*[tiab] OR Pamm1*[tiab] OR maternal serum[tiab] | 161,646 | *Identify relevant monocyte terms or serum studies.* |
| #4 | (#1 OR #2) AND #3 | 2,283 | *Identify relevant combinations of placenta and monocyte studies.* |
| #5 | #4 NOT [Review] | 2,121 | *Exclude review studies.* |
| #6 | Animals[MeSH] NOT Humans[MeSH] | 5,148,047 | *Identify animal studies.* |
| #7 | #5 NOT #6 | 1,768 | *Exclude animal studies.* |
| #8 | Limit to English | 1,634 | *Exclude studies not in English.* |
| #9 | Limit to publishing year: 1995-2023 | 1,193 | *Exclude outdated studies.* |

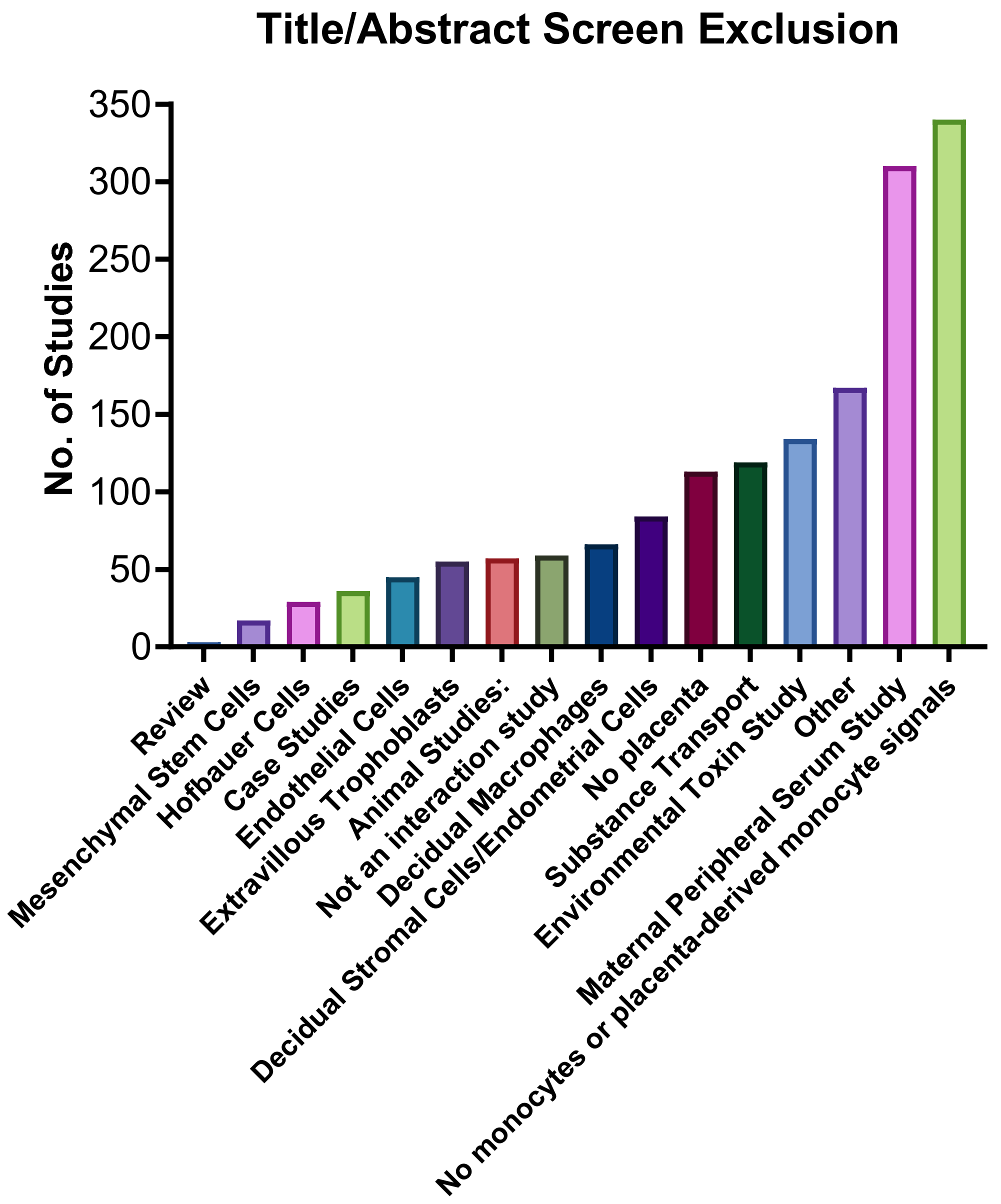


***Supplemental Figure 1: Reasons for article exclusion during the title/abstract screening step.*** *Articles whose title and abstract did not meet the inclusion criteria were excluded and categorized based upon the following categories. Excluded articles were able to be placed into multiple categories.*

**Supplemental Table 2: Data Extraction Questions for Included Articles**

| Category: | Information Extracted: |
|---|---|
| General | • PMID<br>• Year published<br>• Study design<br>• Monocyte/trophoblast interaction studied<br>• Disease State<br>• Disease induction or isolation<br>• Placental tissue source<br>• Mode of delivery<br>• Identification of labored status<br>• Placental tissue type<br>• Placental tissue gestational age<br>• Placental Inclusion Criteria<br>• Study location<br>• Placental tissue control<br>• Monocyte type used<br>• Monocyte source<br>• Peripheral monocyte sex<br>• Pregnancy state of peripheral monocytes<br>• Monocyte control |
| Monocyte Presence/Recruitment | • Experimental methods<br>• Quantification of monocyte numbers<br>• Monocyte identification markers<br>• Relevant changes and p values in disease contexts<br>• Impact of gestational age on monocyte recruitment |
| Monocyte Phenotype in Intervillous Space | • Collection procedure for intervillous blood<br>• Monocyte characterization marker<br>• Method of Detection<br>• Monocyte experimental groups and controls<br>• Relevant marker changes and p values |
| Placental Extracellular Vesicles | • Source of STBEVs<br>• EV types studied<br>• EV isolation methods<br>• Centrifuge speeds<br>• EV characterization methods<br>• Cytokine content of EVs |

| | |
|---|---|
| | • Monocyte characterization markers<br>• Monocyte characterization methods<br>• Relevant changes and P values |
| Monocyte-Placental Adhesion | • Receptors studied<br>• Adhesion ligands studied<br>• ICAM-1 expression changes<br>• CX3CR1 expression changes<br>• Monocyte-trophoblast adhesion changes in disease<br>• Relevant p values |
| Other | • Summarized main findings of the paper. |
| Monocyte Impact on Placenta | • Monocyte effect on trophoblasts |

**Supplemental Table 3: PMIDs of Included Articles and Gestational Ages of Tissue Used**

| Article Title: | PMID: | Article Categories: | Average Gestational Age (GA) of Placenta: |
|---|---|---|---|
| Lysophosphatidic acid up-regulates expression of growth-regulated oncogene-alpha, interleukin-8, and monocyte chemoattractant protein-1 in human first-trimester trophoblasts: possible roles in angiogenesis and immune regulation | 19906815 | Presence | 7-10 weeks gestation |
| Intercellular adhesion molecule-1 expression in massive chronic intervillositis: implications for the invasion of maternal cells into fetal tissues | 24631282 | Presence | 38(34-40) weeks for MCI, 39(34-41) weeks for villitis, 38 (34-40) weeks for non-villitis |
| Plasmodium falciparum infection dysregulates placental autophagy | 31805150 | Presence | 39 (39-40) weeks for control, 39 (38-40) weeks for general malaria infection, 39 (38-40) weeks for malaria infection no placental infection, 39(38-40) weeks for malaria infections with placental infections |
| Massive chronic intervillositis of the placenta associated with malaria infection | 9706981 | Presence | MCI: 37.31 ± 1.59 weeks |
| Maternal malaria induces a procoagulant and antifibrinolytic state that is embryotoxic but responsive to anticoagulant therapy | 22347435 | Presence | Uninfected: 38(2) weeks, placental malaria: 38(1) weeks, placental malaria submicroscopic: 38(1) weeks |
| Placental infection with Plasmodium vivax: a histopathological and molecular study | 23053630 | Presence | Unspecified GA, live deliveries |

| | | | |
|---|---|---|---|
| The Relationship Between Fetal Weight with Sequestration of Infected Erythrocyte, Monocyte Infiltration, and Malaria Pigment Deposition in Placenta of Mother Giving Birth Suffering from Plasmodium Vivax Infection | 34759450 | Presence | Unspecified GA, live deliveries |
| Immunohistopathological changes in the placenta of malaria-infected women in unstable transmission setting of Aligarh | 35970103 | Presence | Unspecified GA, live deliveries |
| Placental blood leukocytes are functional and phenotypically different than peripheral leukocytes during human labor | 19748682 | Presence | 37-40 weeks |
| Comparative flow cytometric analysis of term placental intervillous and peripheral blood from immediate postpartum women in Western kenya | 12852869 | Presence | Term placenta, unspecified GA |
| Human immunodeficiency virus co-infection increases placental parasite density and transplacental malaria transmission in Western Kenya | 19141849 | Presence | Term placenta, unspecified GA |
| Role of some biomarkers in placental malaria in women living in Yaoundé, Cameroon | 25447267 | Presence | Unspecified GA, live deliveries |
| Placental malaria is associated with cell-mediated inflammatory responses with selective absence of natural killer cells | 11237836 | Presence | Unspecified GA, live deliveries |
| Host response to malaria during pregnancy: placental monocyte recruitment is associated with elevated beta chemokine expression | 12594307 | Presence | Unspecified GA, live deliveries |
| Both "canonical" and "immunological" preeclampsia subtypes demonstrate changes in placental immune cell composition | 31477208 | Presence | Selected 5 placentas from each “cluster” of previously published set of data. Unclear the gestational ages of the samples, but the overall groups: Cluster 1 36(4) weeks, Cluster 2 31(3) weeks and Cluster 3 34(4) weeks[1] |
| Diffuse and Localized SARS-CoV-2 Placentitis: Prevalence and Pathogenesis of an Uncommon Complication of COVID-19 Infection During Pregnancy | 35319524 | Presence | 7 Cases were: 27.7, 37.0, 32.7, 29.1, 38.7, 40.6, 35.1 weeks at delivery |
| Significance of C4d Immunostaining in Placental Chronic Intervillositis | 25970733 | Presence | 13-40 weeks gestation, list of individual sample records in text |
| Expression analysis of leukocytes attracting cytokines in chronic histiocytic intervillositis of the placenta | 23696928 | Presence | CHI: 27 (19-33) weeks, VUE 37(35-39) weeks, Control 27 (20-33), 37(36- |

| | | | |
|---|---|---|---|
| | | | 38) weeks (age-matched the control) |
| Activation of leukocytes during the uteroplacental passage in preeclampsia | 11799095 | Phenotype | No placental tissue collected. GA of uterine vein blood: Normal: 38.3 (38.2-38.4) weeks, Preeclampsia: 30.6(29.2-32.4) weeks |
| IL-12 producing monocytes and IFN-gamma and TNF-alpha producing T-lymphocytes are increased in placentas infected by Plasmodium falciparum | 17194481 | Phenotype | Term Placenta, Unspecified GA |
| Monocyte activation and T cell inhibition in Plasmodium falciparum-infected placenta | 15181571 | Phenotype | Unspecified GA, live deliveries |
| Placental fractalkine mediates adhesion of THP-1 monocytes to villous trophoblast | 25566740 | Adhesion | Cell line |
| Macrophage migration inhibitory factor is up-regulated in human first-trimester placenta stimulated by soluble antigen of Toxoplasma gondii, resulting in increased monocyte adhesion on villous explants | 18165264 | Adhesion | 9-12 weeks gestation |
| Toxoplasma gondii regulates ICAM-1 mediated monocyte adhesion to trophoblasts | 16174097 | Adhesion | Cell line |
| ICAM-1-mediated adhesion of peripheral blood monocytes to the maternal surface of placental syncytiotrophoblasts: implications for placental villitis | 9137107 | Adhesion | Term placenta, unspecified GA |
| Human placental exosomes induce maternal systemic immune tolerance by reprogramming circulating monocytes | 35180876 | EVs | 7-10 weeks gestation |
| Systemic inflammatory priming in normal pregnancy and preeclampsia: the role of circulating syncytiotrophoblast microparticles | 17442979 | EVs | Term placenta, 39-41 weeks gestation |
| The immunomodulatory role of syncytiotrophoblast microvesicles | 21633494 | EVs | 37.3 (35.3 to 39.6) weeks |
| A First Phenotypic and Functional Characterization of Placental Extracellular Vesicles from Women with Multiple Sclerosis | 33809077 | EVs | MS: 37 (2.7) weeks, Healthy control: 38.2 (1.7) weeks |
| Feto-maternal interactions in pregnancies: placental microparticles activate peripheral blood monocytes | 20005571 | EVs | Term placenta, unspecified GA |
| Trophoblast cells are able to regulate monocyte activity to control Toxoplasma gondii infection | 23294571 | Other | Cell line |

| | | | |
|---|---|---|---|
| BEWO trophoblast cells and Toxoplasma gondii infection modulate cell death mechanisms in THP-1 monocyte cells by interference in the expression of death receptor and intracellular proteins | 34597888 | Other | Cell line |
| Natural hemozoin stimulates syncytiotrophoblast to secrete chemokines and recruit peripheral blood mononuclear cells | 21632106 | Other | Term placenta, unspecified GA |
| Interleukin-34 is present at the fetal-maternal interface and induces immunoregulatory macrophages of a decidual phenotype in vitro | 29579271 | Other | Elective termination: 9(6-13) weeks, Early-onset PE <32 weeks, Normal term pregnancies and late-onset PE samples, unspecified GA |
| Inflammasome signaling in human placental trophoblasts regulates immune defense against Listeria monocytogenes infection | 32976558 | Other | Term for C-section placentas, unspecified GA. <24 weeks gestation for tissue from elective terminations |
| Plasmodium falciparum malaria elicits inflammatory responses that dysregulate placental amino acid transport | 23408887 | Mono Impact | Cell line |
| Macrophage- but not monocyte-derived extracellular vesicles induce placental pro-inflammatory responses | 30213492 | Mono Impact | Term placenta, unspecified GA |
| Enhanced monocyte binding to human cytomegalovirus-infected syncytiotrophoblast results in increased apoptosis via the release of tumour necrosis factor alpha | 16158462 | Adhesion, Mono Impact | Term placenta, unspecified GA |
| Monocytes adhering by LFA-1 to placental syncytiotrophoblasts induce local apoptosis via release of TNF-alpha. A model for hematogenous initiation of placental inflammations | 11129659 | Adhesion, Mono Impact | Term placenta, unspecified GA |
| Phenotypic and functional characterization of first-trimester human placental macrophages, Hofbauer cells | 33075123 | Adhesion, Phenotype, Presence | 6-12 weeks gestation |
| Expression of intercellular adhesion molecule 1 (ICAM-1) in Plasmodium falciparum-infected placenta | 11440546 | Adhesion, Presence, Phenotype | Uninfected:36.6(0.9) weeks, Acute placental malaria: 35.5 (2.6) weeks, chronic placental malaria: 35.4(2.5) weeks, Overall placental malaria: 35.5(2.5) weeks |
| Maternal Monocytes Respond to Cell-Free Fetal DNA and Initiate Key Processes of Human Parturition | 34663619 | Other | Term placenta (unspecified GA range) |

| | | | |
|---|---|---|---|
| The effect of kisspeptin on the functional activity of peripheral blood monocytes and neutrophils in the context of physiological pregnancy | 35413647 | Other | None |
| Labour promotes systemic mobilisation of monocytes, T cell activation and local secretion of chemotactic factors in the intervillous space of the placenta | 36969250 | Phenotype, Presence | 39 (39-40) weeks for non-labor, 41 (39-41) weeks for labor |
| Changes in the levels of chemokines and cytokines in the placentas of women with Plasmodium falciparum malaria | 14513430 | Phenotype, Presence | Term placenta, unspecified GA |
| ICAM-1 expression on immune cells in chronic villitis | 25454473 | Presence, Adhesion | Term/near term placentas, unspecified GA |
| The activating effect of IFN-γ on monocytes/macrophages is regulated by the LIF-trophoblast-IL-10 axis via Stat1 inhibition and Stat3 activation | 25027966 | Presence, Other | Cell line |
